# *In silico* analysis and molecular investigation of the nuclear shuttle protein-interacting kinase 1 (NIK1) in six Solanaceous species

**DOI:** 10.1101/2021.07.06.451341

**Authors:** Mehdi Safaeizadeh, Nachelli Malpica

## Abstract

The first layer of innate immunity in plants is initiated through the perception of microbe-associated molecular patterns (MAMPs) or damage-associated molecular patterns (DAMPs) by pattern recognition receptors (PRRs). MAMP/DAMP perception initiates downstream defense responses, a process which ultimately leads to pattern triggered immunity (PTI). In Arabidopsis, the nuclear shuttle protein-interacting kinase 1 (NIK1), among other PRRs, is one of the most important central components of PTI signaling and kinase signaling cascade, since it is involved in the plant antiviral response against geminiviruses. Despite the characterization of the structure and function of the NIK1 receptors made by some groups, studies related to NIK1 importance in the current gene-editing era are missing. By simple *in silico* analysis, in this study we investigated the NIK1 homologues from six Solanaceous plant species including: tomato (*Solanum lycopersicum*), potato (*Solanum tuberosum*), *Solanum pennellii*, eggplant (*Solanum melongena*), pepper (*Capsicum annum*), and *Nicotiana benthamiana*. The phylogenetic analyses of different NIK1 proteins from Arabidopsis and six Solanaceous plants revealed nine different clades. As expected, we found that these NIK1 orthologs have similar genomic structures suggesting a similar function. We could identify that SotubNIK1, SolyNIK1, SopenNIK1, CANIK1, NibenNIK1, SmeNIK1 have the highest sequence homology with AtNIK1. Additionally, the conserved protein kinase domain (PKD) that is present in NIK1 from *Arabidopsis thaliana* was bioinformatically analyzed and found in other species. As this highly conserved NIK1 region is present in several crops of economic importance, its potential is highlighted as a possible target site for gene editing, to develop crops tolerant to geminiviral infections.

## Introduction

Plants and animals are always at risk of infections from various micro-organisms. There are several observed similarities between the plant and animal immune systems^1,2^, but as plants lack specialized mobile immune cells, as well as the adaptive immune system that is present in animals, they largely depend on their innate immune system to defend themselves against potential infections^3^.

Plants are equipped with different arrays of defense strategies^4^. In a co-evolutionary arm race between plants and pathogen interactions, as a first line of defense against microbial pathogens, plants perceive microbes via microbe-associated molecular patterns (MAMPs) by plasma membrane-resident immune receptors which are called pattern-recognition receptors (PRRs). These receptors are located on the cell surface^4,5^. As a consequence of MAMPs perception, a signaling cascade is initiated that finally leads to pattern-triggered immunity PTI^4^, where global transcriptional changes and cell re-programming take place^8^. Interference with this network can paralyze the adequate response upon pathogen infection^6-7^.

Besides PTI, there is another layer of the immune system called effector triggered immunity (ETI), which is initiated by the activation of nucleotide-binding leucine-rich repeat (NLR) proteins. These receptors detect the presence of pathogen-derived effectors within plant cells, but the underlying mechanisms of recognition are still poorly understood^9^. Overlapping responses between PTI and ETI have been observed, although ETI has been shown to be stronger than PTI leading to plant cell death. Like the plant innate immune system shares a common ancestor, the better understanding of the defense mechanisms in the model plant *Arabidopsis thaliana* offers an opportunity to formulate adequate defense strategies against crop diseases^10^.

*A. thaliana* is able to detect the presence of a variety of MAMPs including bacterial elicitors like flagellin^11^, the elongation factor EF-Tu^11^, peptidoglycans^12^, lipopolysaccharides^13^, bacterial cold shock proteins^4^, bacterial superoxide dismutase^13^, conserved bacterial peptides^4^, bacterial outer membrane vesicles^14^ and more recently bacterial RNAs^15-16^ However, despite the identification of some of these elicitor receptors ^17^, many of them remain still unknown.

Plant PRRs are subdivided by the nature of their ligand-binding ectodomain^1^ and include receptor-like kinases (RLKs) and receptor-like proteins (RLPs) which lack a cytoplasmic kinase domain. RLKs which are involved in ligand binding, possess a variable extracellular domain, a single transmembrane domain and an intracellular kinase domain^1^. Genome-wide investigation in *Arabidopsis thaliana* showed that RLKs are part of the gene family that shares a common ancestor with the animal receptor tyrosine kinases (RTKs), and receptor serine and threonine kinases (RSKs). Data has shown that more than 600 RLKs constitute 2.5% of all the protein-coding genes in Arabidopsis^19a^. The motifs making up the extracellular part of RLKs vary greatly among members, the most common being leucine-rich repeats (LRRs) (235 of the 610 RLKs contain 1–32 LRRs in their extracellular domain) ^19a^. LRR-RLKs have a role in many different important signaling processes such as immunity^20^, hormone signaling^21^, and development^22^. The structural plasticity of LRR-containing proteins allows them to bind to many distinct ligands, including peptides, lipids, hormones and other biochemical compounds^1^

The family of LRR-RLKs has been divided into 13 groups (LRR-RLK I to LRR-RLK XIII)^19b^. Recent phylogenetic studies showed that the somatic embryogenesis receptor kinases (SERKs) together with NIK1 and NIK2 receptors, belong to clade 1 of the five clades described for the LRR-RLK II group of receptor-like kinases in the plant kingdom^35^. The SERK family, including SERK1, SERK2, BAK1/SERK3, and BAK1-LIKE1/SERK4 (BKK1/SERK4) is involved in plant development and disease resistance and they serve as co-receptors in brassinosteroids and immune signaling (Li et al., 2002, Nam and Li, 2002, Chinchilla et al., 2007). The SERK proteins have a small extracellular domain consisting of 4.5–5 LRRs. They distinguish themselves from other LRR-RLK II members by the presence of a serine-proline rich (SPP) motif in their extracellular domain^23^. Phylogenetic analysis of SERK proteins in *A. thaliana* and tomato indicates that SERKs constitute a highly conserved protein family^24-25^. Despite a partial redundancy in their functions, each member of the SERK family performs a very specific subset of signaling roles and each member has a specific function^24-25^. The functional specificity of this highly homologous protein family is still not fully understood, and there are many questions regarding the evolution and functional properties of many of its members^24^. The short amino acid motifs present in their extracellular domains are highly conserved^24-25^ and these are important for their interaction with other receptor kinases^24-25^. BAK1/SERK3, one of the best characterized members of this family, is known to act as a coreceptor of the Brassinosteroid insensitive (BRI-1) receptor, involved in cell elongation and tolerance to environmental stress, and of other several PRRs related to innate immunity like FLS2, PEPR1, PEPR2 and EF-TU receptor (EFR)^27^.

In Arabidopsis the nuclear shuttle protein (NSP)-interacting kinase 1 (NIK1) has been identified as an LRR-RLK II receptor involved in the recognition of geminiviruses. One of its most important features is the phosphorylation of the threonine (T) residue 474 present in its kinase domain that is fundamental for its activation^25 36^ to promote the inhibition of the viral and host mRNA translation. The model proposes that the phosphorylation of threonine leads to the phosphorylation of the downstream component ribosomal protein L10A (RPL10), which interacts in the nucleus with the L10-interacting MYB domain-containing protein (LIMYB), to down regulate genes associated with the translational machinery of the host. This one also leads to the translational inhibition of the virus and an increased tolerance to geminiviruses by the infected plants^37,37b^. As a counter-defense, the viral nuclear shuttle protein (NSP) binds to the kinase domain of NIK1 to inhibit T474 phosphorylation and activation, favoring viral infection. Interestingly, in recent years Santos et al., 2009, reported a gain-of-function mutation in the NIK1 kinase domain (T474D) from Arabidopsis that leads to the hyperactivation of NIK1 kinase activity and resistance against geminiviral infection when expressed in tomato (Brustolini et al.,2015).

Since viral diseases cause important economical loses in solanaceous plants^27-28^ and other important crops like beans, manioc and maize, the identification of the genes which can have a role in antiviral defense in plants becomes necessary. Despite the extensive research done on signaling receptors and proven participation of NIK1 in signaling cascades against viral infection, there is few published data on the phylogenetic analysis of NIK1 in solanaceous plants and in other crops.

In the current publication, the phylogenetic analysis of NIK1 at the amino-acid level in *Arabidopsis thaliana* and six important solanaceous plants is shown. After performing multiple sequence alignment, domain analysis and protein structure analysis, we could identify six NIK1 orthologs in the following solanaceous plants including tomato (*Solanum lycopersicum*), potato (*Solanum tuberosum*), *Solanum pennellii*, eggplant (*Solanum melongena*), pepper (*Capsicum annum*), and *Nicotiana benthamiana*. Additionally, the NIK1 kinase domain was also identified in other important crops. The present research could be the base to characterize the function of NIK1 orthologs in Solanaceous plants and other crops of economic importance.

## Materials and methods

### Database searches

To identify NIK1-related proteins in the six solanaceous species, the amino acid sequence of AT5G16000 (NIK1) in *Arabidopsis thaliana* was directly obtained from TAIR (https://www.arabidopsis.org/) and used as query to blast against the six solanaceous species using the tBLASTN program in the Sol Genomics Network (SGN) (https://solgenomics.net), TAIR (https://www.arabidopsis.org/), Phytozome (columbine), and the National Center for Biotechnology Information (NCBI) (https://www.ncbi.nlm.nih.gov/) websites to identify possible NIK1 orthologs. All searches were cross analyzed to compile a list of the best possible candidates. Protein sequences with more than 90% sequence identity were considered.

### Multiple sequence alignment analysis

Multiple alignment analysis of the sequences was performed based on the Clustal Omega software^93^.The protein sequence alignment of NIK1 from *Arabidopsis thaliana*, tomato (*Solanum lycopersicum*), potato (*Solanum tuberosum*), *Solanum pennellii*, eggplant (*Solanum melongena*), pepper (*Capsicum annum*), and *Nicotiana benthamiana* was done, and the alignments were manually corrected and trimmed to obtain comparable sequences (Figure 1). The protein sequence alignment of Arabidopsis NIK1 kinase domain was also done using the same software (Figure 6).

**Figure 1.**
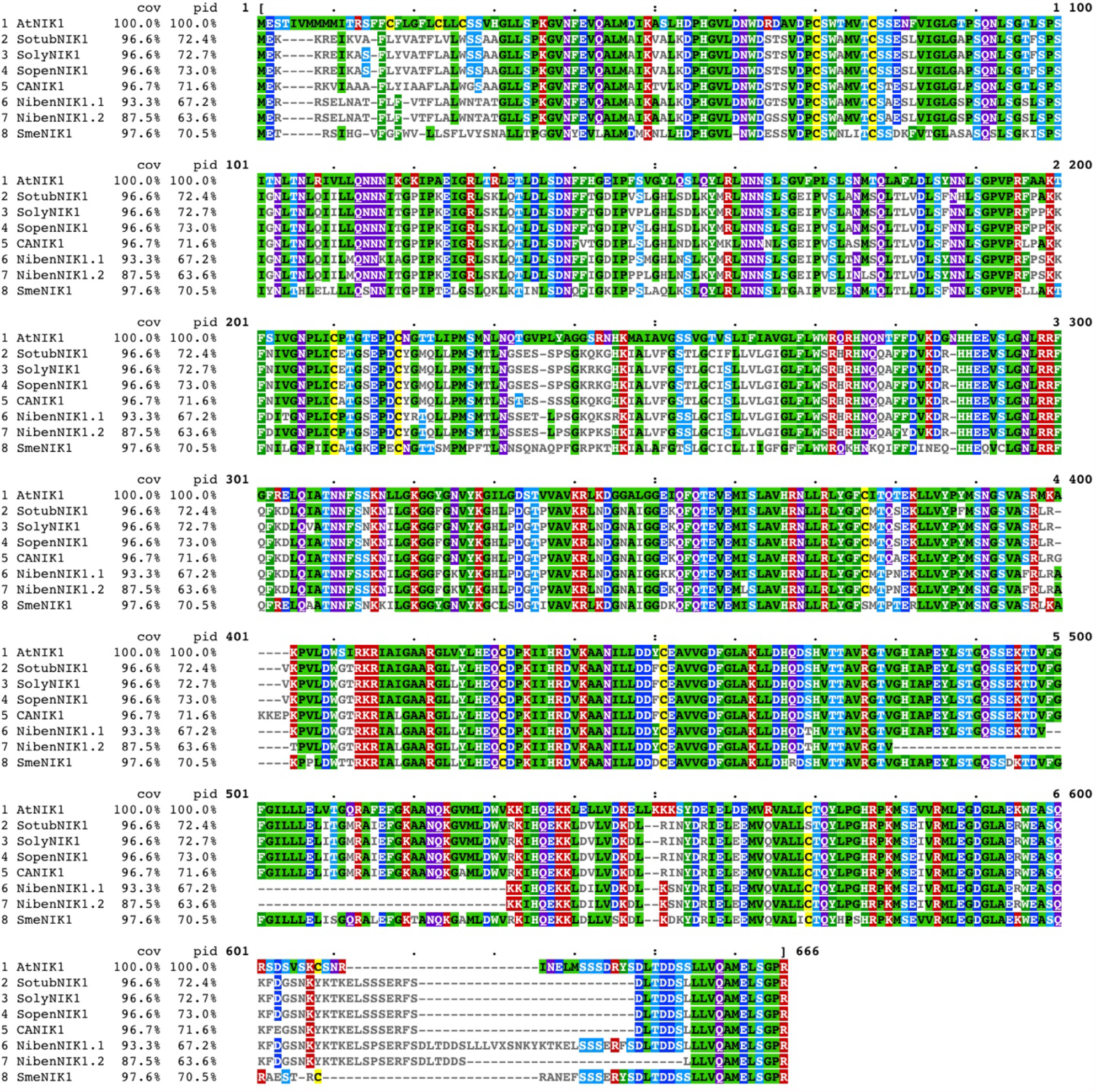
Multiple amino acid sequence alignment of NIK1 in *Arabidopsis thaliana* and six Solanaceous species including: AtNIK1 in *A. thaliana;* SopenNIK1 in *Solanum tuberosum*; SolyNIK1 in *Solanum lycopersicum*; SopenNIK1 in *Solanum pennellii;* CANIK1 in *Capsicum annum*; SmeNIK1 in *Solanum melongena*; NibenNIK1.1 and NibenNIK1.2 in *Nicotiana benthamiana*. The alignment was carried out by using Clustral Omega.

### Phylogenetic analyses and comparison of the amino acid sequences

The phylogenetic analysis was carried out using MEGA software (Molecular Evolutionary Genetics Analysis) version X (Kumar *et al*., 2018). A maximum likelihood phylogenetic tree was constructed using default parameters and 1000 bootstrap replicates. All branches with bootstrap values < 60 were collapsed.

### Structure analysis of NIK1 amino acid sequences and motif investigation

Using the NPS@SOPMA (https://npsa-prabi.ibcp.fr/) with its default settings, the secondary structures of NIK1 proteins including AtNIK1, SotubNIK1, SolyNIK1, SopenNIK1, CANIK1, NibenNIK1, SmeNIK1 were predicted. The investigation of the structure analysis included the percentage of each amino acid and position and the alpha helix number. In addition, βeta bridges and random coils were identified. To determine the motifs in the NIK1 amino acid sequences, motif analysis was performed using the MEME tool with motif number set as 10. The other parameters were set as default. The Swiss-Model (https://www.swissmodel.expasy.org/), which is based on the automatic ExPASy (Expert Protein Analysis System) (Waterhouse et al., 2018; Waterhouse et al., 2016), was used to determine the 3D models of the protein tertiary structures of AtNIK1, SotubNIK1, SolyNIK1, SopenNIK1, CANIK1, NibenNIK1, SmeNIK1.

## Results

### Identification and phylogenetic analysis of the NIK1 proteins in six Solanaceous plants

Taking the NIK1 gene AT5G16000.1 from *Arabidopsis thaliana* (AtNIK1) as a reference, we could identify a total of 66 NIK1 homologs with top hits in the six studied solanaceous plants including: 11 from *Arabidopsis thaliana*, 10 from potato (*Solanum tuberosum*), 10 from tomato (*Solanum lycopersicum*), nine from *Solanum pennellii*, six from eggplant (*Solanum melongena*), seven from pepper (*Capsicum annum*), and 13 from *Nicotiana benthamiana*. (Supplementary Dataset S1). All the proteins showed the highest sequence similarity with AtNIK1. The amino acid sequences among the solanaceous plants were highly conserved, suggesting that NIK1 orthologs exist in Solanaceous plants. To better observe the relatedness among AtNIK1 and its possible orthologs in *Solanaceae*, an un-rooted phylogenetic tree was constructed with the full-length sequences of the selected proteins through a maximum likelihood phylogenetic analysis. (Supplementary Dataset S2). The found homologues were named according to the corresponding species as shown in Figure 2.

**Figure 2.**
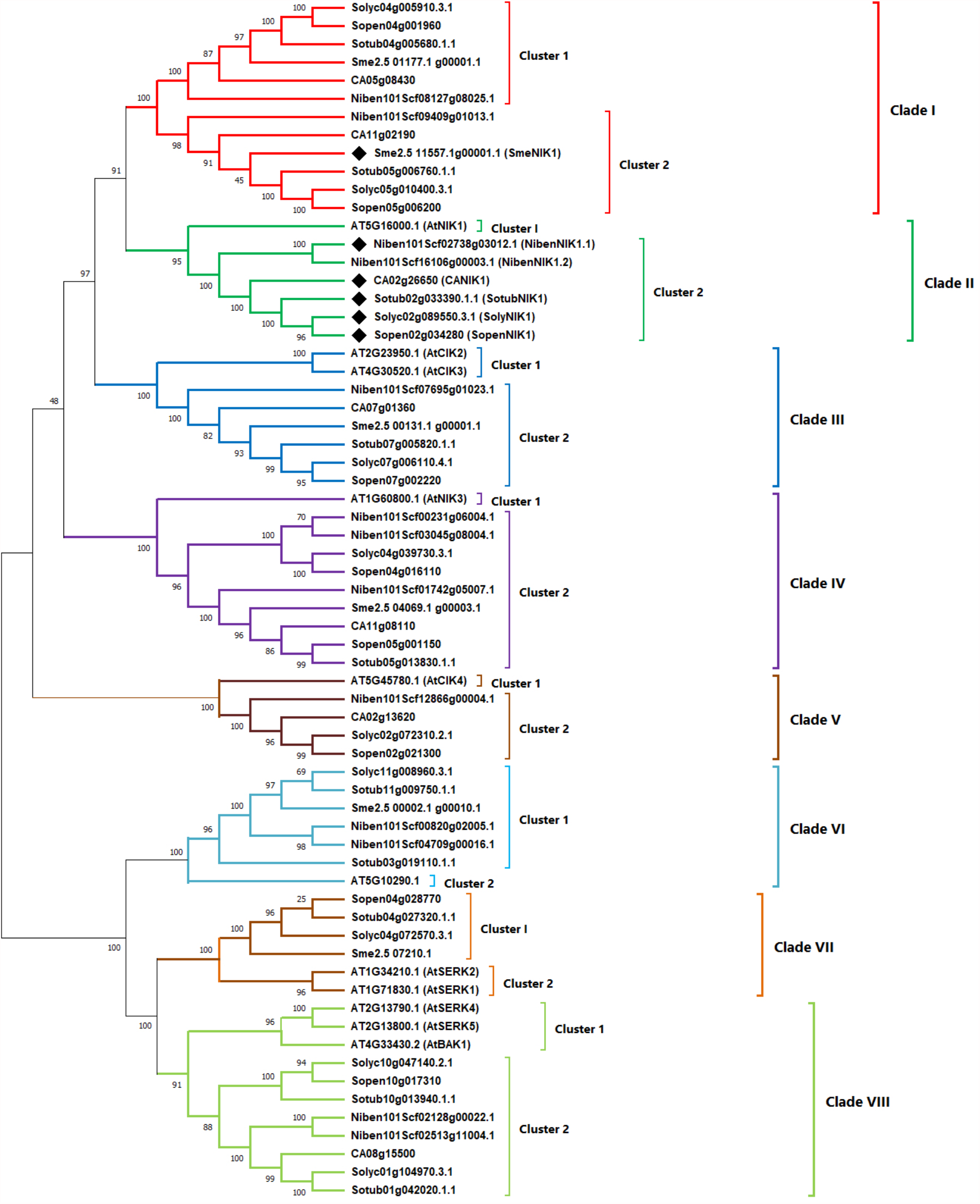
Phylogenetic analysis of the *Arabidopsis thaliana* AtNIK1 protein sequence and its logs in six representative solanaceous plants. The species indicated are potato (*Solanum osum*), tomato (*Solanum lycopersicum*), *Solanum pennellii*, eggplant (*Solanum melongena*), er (*Capsicum annum*), and *Nicotiana benthamiana* protein sequences homologs. AtNIK1 protein *abidopsis thaliana* was labelled. Sequences for comparisons were obtained from Solgenomic.net ank. The accession numbers and protein names (if available) are given. Analysis was done by mum likelihood method with 500 bootstrap replicates implemented in MEGAX (Molecular utionary Genetics Analysis) version X.

**Figure 3.**
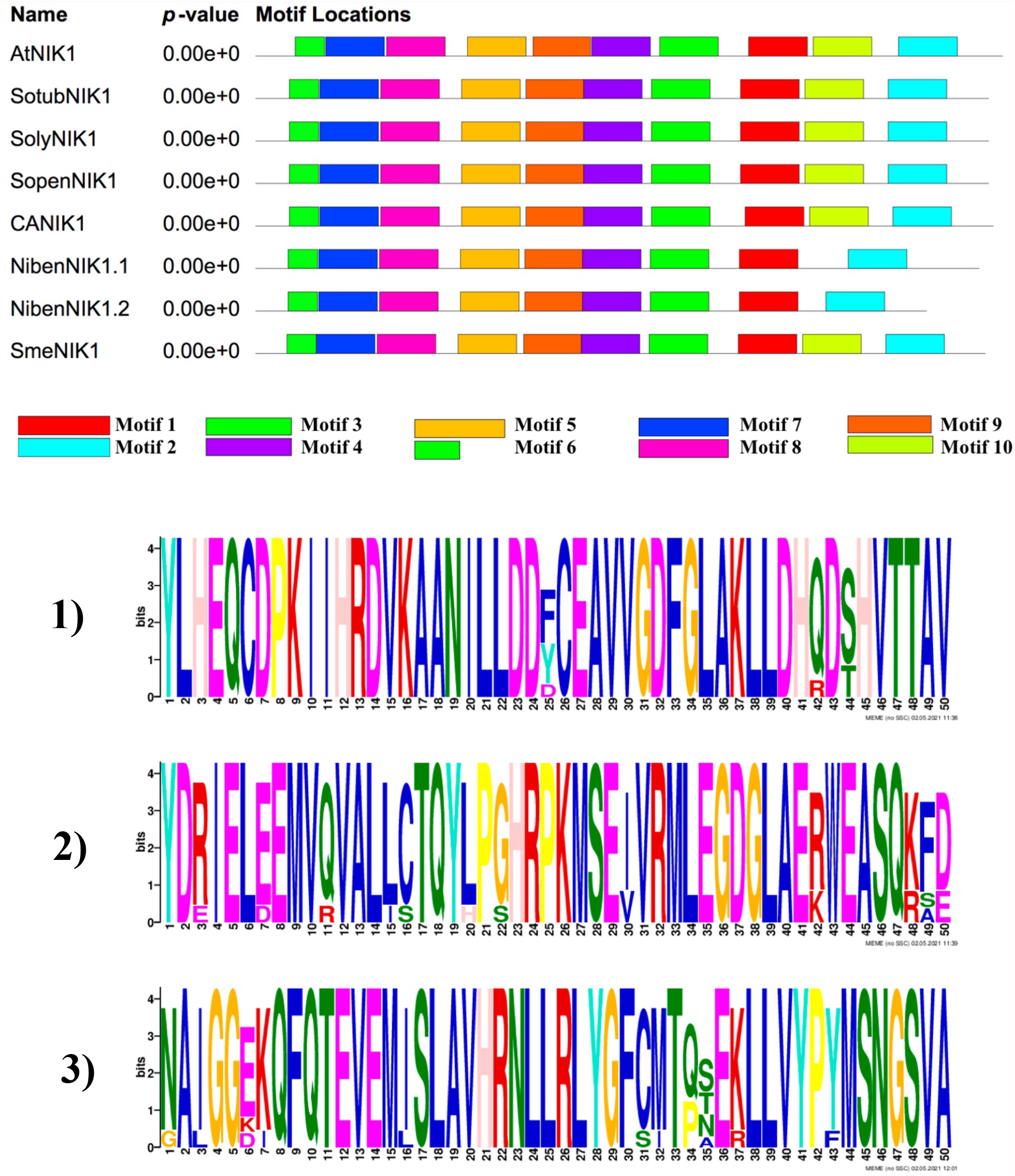
Conserved protein motif analysis of NIK1 in *Arabidopsis thaliana* and six Solanaceous species including: AtNIK1 in *A. thaliana;* SopenNIK1 in *Solanum tuberosum*; SolyNIK1 in *Solanum lycopersicum*; SopenNIK1 in *Solanum pennellii;* CANIK1 in *Capsicum annum*; SmeNIK1 in *Solanum melongena*; NibenNIK1.1 and Niben NIK1.2 in *Nicotiana benthamiana*. Conserved motifs in the NIK1 proteins were elucidated by MEME website (version 5.0.3). 10 motifs were shown by different colors. Different motifs were represented in different colors in the protein. The details of the conserved motifs 1, 2 and 3 which have the highest amino-acid sequence similarity are shown. The height of the amino acids present in each motif is shown in bits.

**Figure 4.**
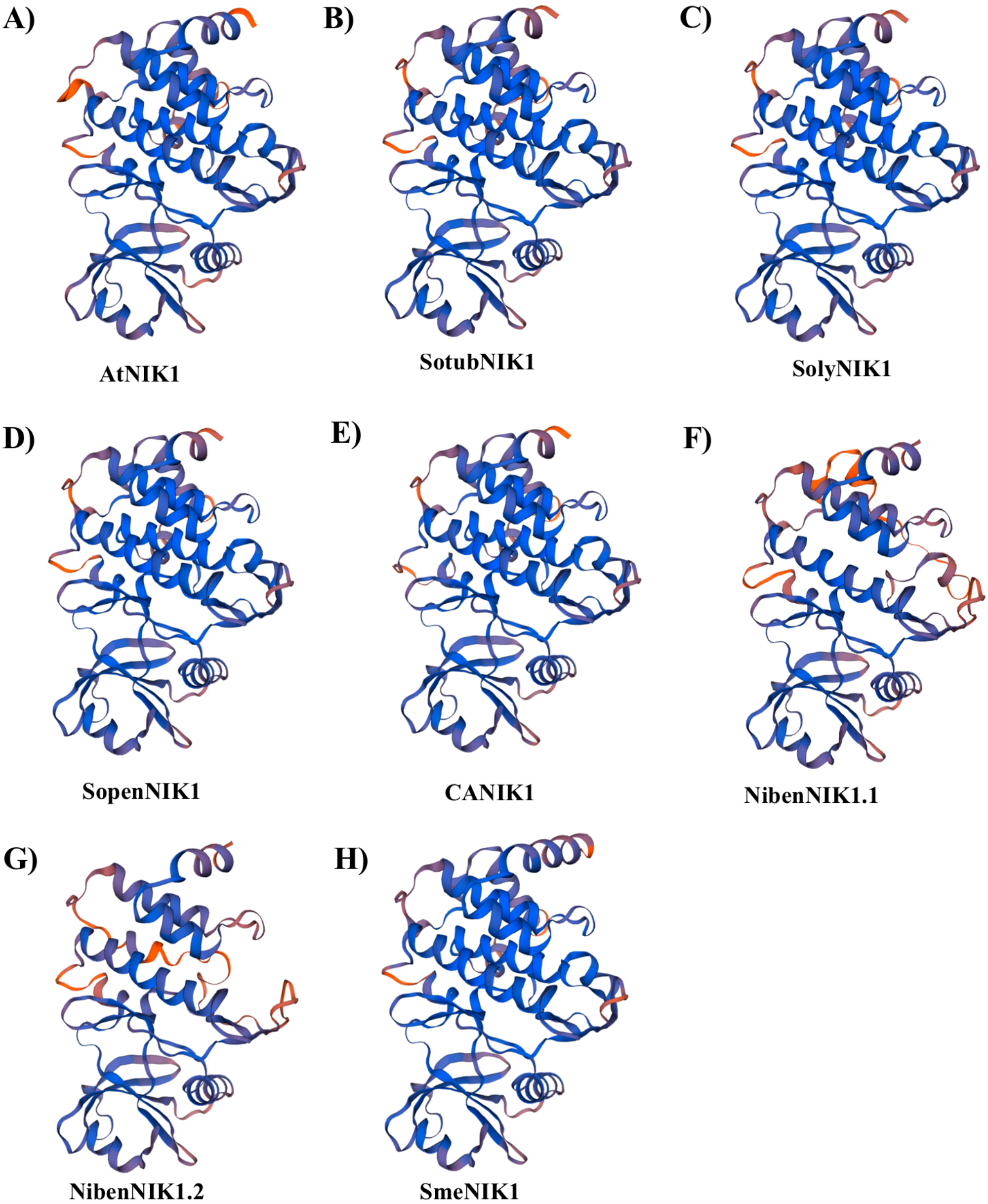
3D structure analysis of NIK1 in *Arabidopsis thaliana* and six Solanaceous species including: AtNIK1 in *A. thaliana;* SopenNIK1 in *Solanum tuberosum*; SolyNIK1 in *Solanum lycopersicum*; SopenNIK1 in *Solanum pennellii;* CANIK1 in *Capsicum annum*; SmeNIK1 in *Solanum melongena*; NibenNIK1.1 and NibenNIK1.2 in *Nicotiana benthamiana*. The structure models were constructed using the Swiss-model.

**Figure 5.**
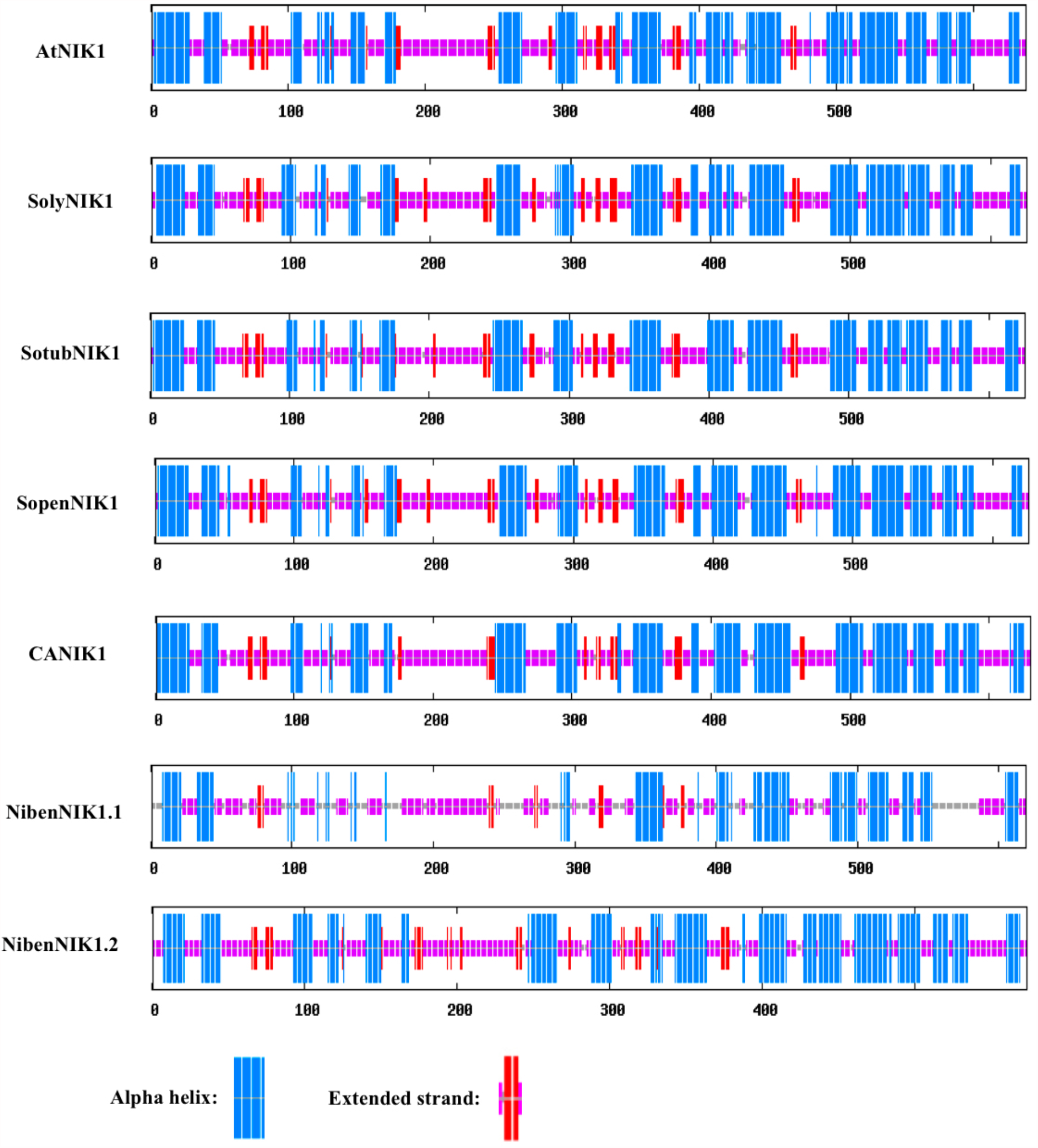
Secondary structure analysis of the NIK1 *A. thaliana* and six Solanaceous plants. AtNIK1 in *A. thaliana;* SopenNIK1 in *Solanum tuberosum*; SolyNIK1 in *Solanum lycopersicum*; SopenNIK1 in *Solanum pennellii;* CANIK1 in *Capsicum annum*; SmeNIK1 in *Solanum melongena*; NibenNIK1.1 and NibenNIK1.2 in *Nicotiana benthamiana*. Alpha helix is colored in blue, extended strand is colored in red. Random coil is colored in purple.

**Figure 6.**
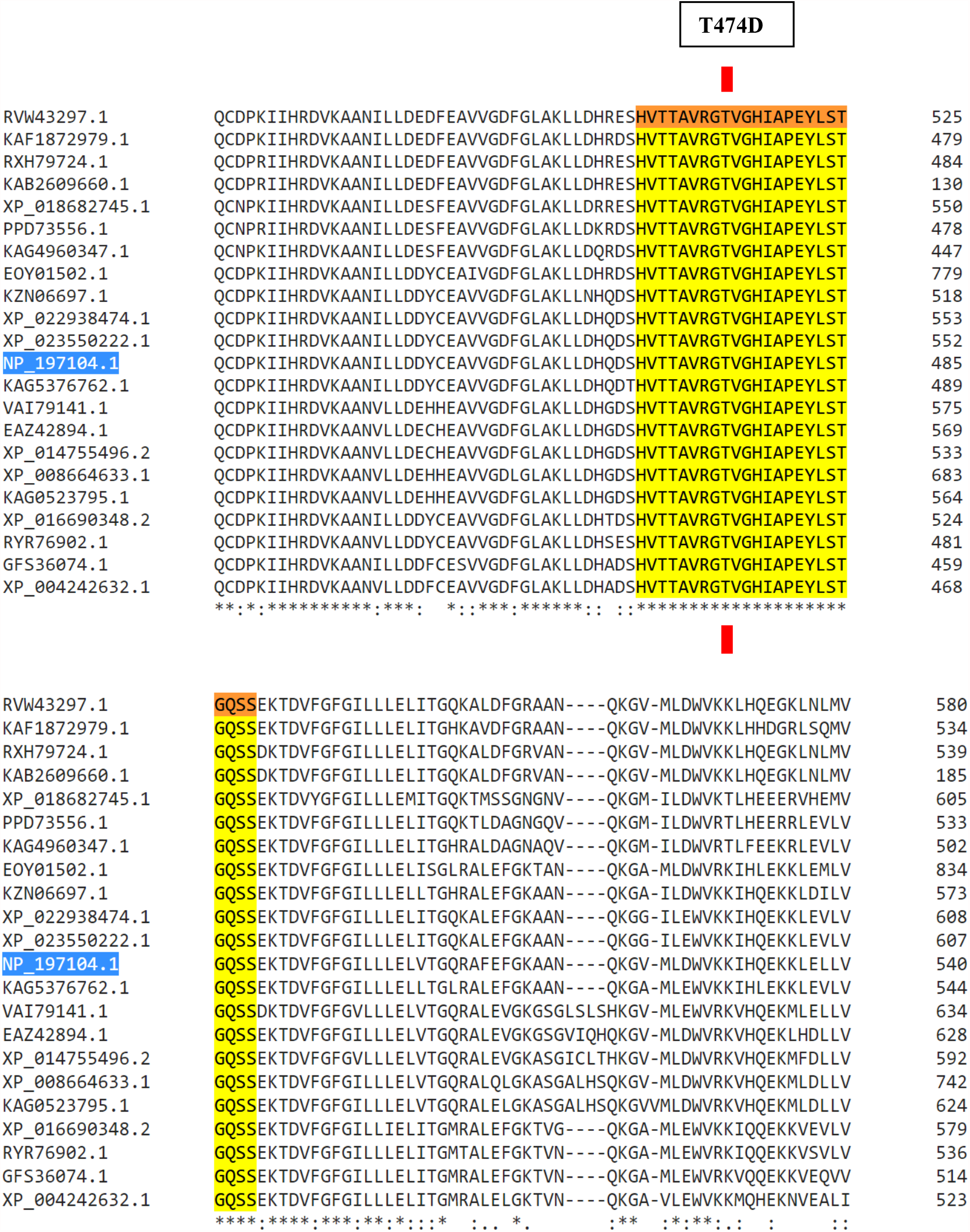
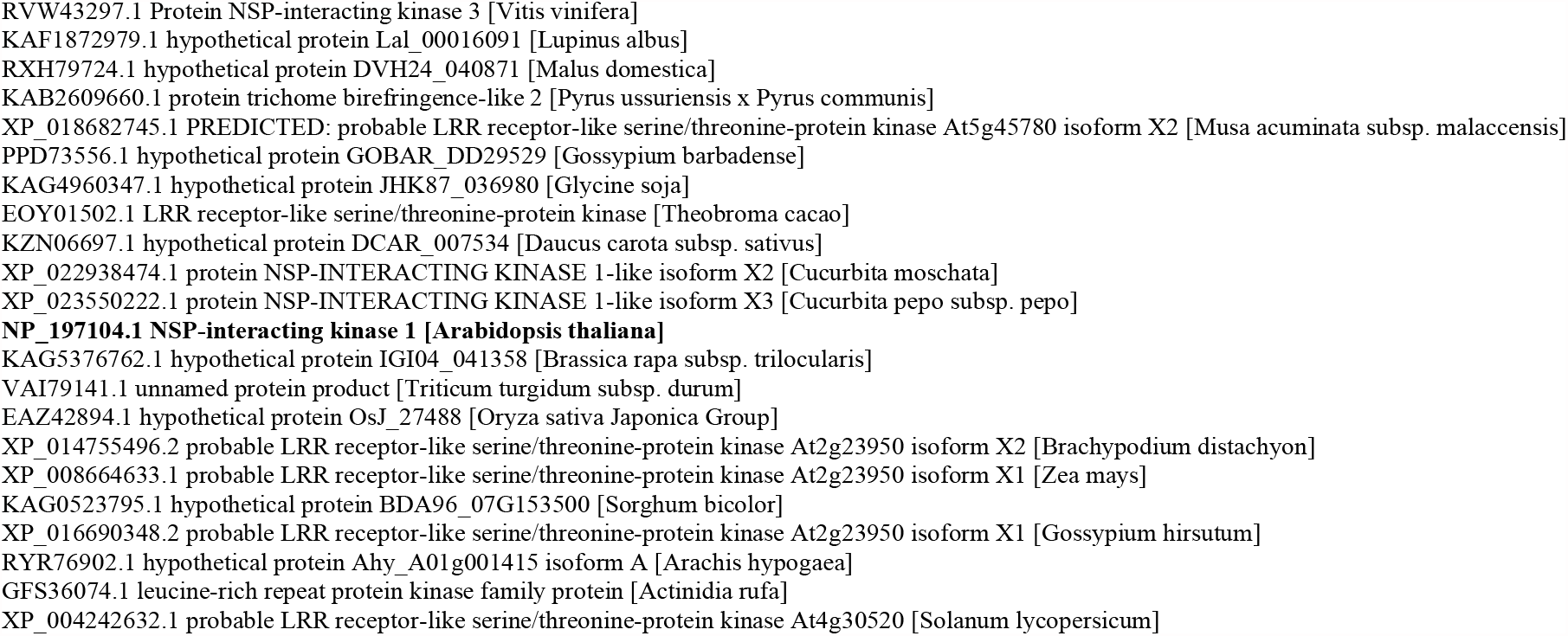
Multiple alignment of the *Arabidopsis thaliana* NIK1 (AT5G16000.1**/**NP_197104.1) kinase domain with other plant species, which could be mutated (T474D) to gain resistance against geminiviral infection. The T474 residue is indicated by the red rectangle on the yellow region showing a section of the kinase domain present in these proteins.

To evaluate and investigate the phylogenetic relationship among them, phylogenetic analysis was investigated in depth. As illustrated in the Figure 2, phylogenetic analysis with bootstrap values of 0 to 100 showed that 66 NIK1-related proteins were found to be divided into eight different clades (Clade I to VIII). A glance at the phylogenetic tree shows that the highly similar sequences with AtNIK1 are present in clades I and II, being clade I the largest clade. Clade I is subdivided in two clusters, each of six sequences. Clade I is comprised of 12 sequences including: two from *A. thaliana*, two from *S. tuberosum*, two from *S. lycopersicum*, two from *S. pennellii*, two from *S. melongena*, two from *C. annum*, and two from *N. benthamiana*.

Clade II is composed of 7 sequences subdivided in two clusters where, cluster 1 contains the NIK1 sequence from *A. thaliana*, and cluster 2 the sequences from *Nicotiana benthamiana*, potato, tomato and chili pepper (two from *N. benthamiana*, one from *S. tuberosum*, one from *S. lycopersicum*, one from *S. pennellii*, and one from *C. annum*).

Clade III consists of eight members and has two clusters. The first cluster includes AtCIK2 and AtCIK3, while the second cluster contains one sequence from *S. tuberosum*, one from *S. lycopersicum*, one from *S. melongena*, one from *S. penellii*, one from *C. annum*, and one from *N. benthamiana*.

The fourth clade has 10 members. Cluster 1 includes one sequence from *A. thaliana* (AtNIK3) and the rest of the nine sequences, which belong to cluster 2, include one sequence from *S. tuberosum*, one from *S. lycopersicum*, one from *S. melongena*, two from *S. penellii*, one from *C. annum*, and three from *N. benthamiana*.

Clade V has five sequences. While AtCIK4 is represented in one cluster, four other members of this clade, comprising the second cluster, include one sequence from *S. lycopersicum*, one from *S. pennellii*, one from *C. annum*, and one from *N. benthamiana*.

Clade VI consists of 7 sequences. The first cluster comprises two sequences from *S. tuberosum*, one from *S. lycopersicum*, one from *S. melongena*, two from *N. benthamiana*, and cluster 2 comprises one sequence from *A. thaliana*.

Clade VII is represented by two clusters. The first cluster consists of four sequences including one from *S. tuberosum*, one from *S. lycopersicum*, one from *S. melongena*, and one from *S. pennellii*. Two members from *A. thaliana* in cluster 2 are related to the SERK1 and SERK2 families.

Clade VIII consists of two clusters. The first one comprising sequences from the SERK4 and SERK5 families together with BAK1, and a second cluster comprising two sequences from *S. lycopersicum*, two from *S. tuberosum*, one from *S. pennellii*, two from N. benthamiana and one from *C. annum*.

### Identification of NIK1 kinase domain in other important species

The results of the multiple alignment of the kinase domain present in *Arabidopsis thaliana* NIK1 also showed that this domain is present in related proteins from other important crops like soy, maize, lupin, pumpkin, rice, cotton, etc. which are also susceptible to geminiviral infection (Figure 6).

## Discussion

In innate immunity, co-receptors play an important role in the activation of RLK-mediated signal transduction cascades. NIK1 is regarded as one of the most important co-receptors and its role in antiviral defense has been confirmed^26^. As solanaceous plants are very sensitive to viral infections including those induced by geminiviruses, a comprehensive analysis of NIK1 orthologs and their biological characterization are necessary, to elaborate possible defense strategies against geminiviral infections. The NIK1 orthologs shown in the present work offer an opportunity to do so. For example, CRISPR/Cas9 has already been used to develop resistance against *Turnip mosaic virus* (TuMV) in *A. thaliana* and resistance in cucumber against other viruses by disrupting the function of translation factors (Pyott et al. 2016; Chandrasekaran et al. 2016). Using the same technology, the introduction of a mutation in the NIK1 kinase domain as shown by Brustolini et. al. (2015) could be an alternative to be considered to develop resistance in solanaceous plants and other important crops like maize, rice and cotton, which also contain similar receptors with the same kinase domain. The present study based on the phylogenetic and structural analysis of NIK1 and its orthologs in other species, had the goal to start to pave the way to study NIK1 receptors with the aim to characterize their functions in *A. thaliana* and solanaceous plants, through experiments that involve NIK1 protein expression or the use of NIK1 mutant plants in combination with infection trials performed with geminiviruses.

## References

1. Couto D, and Zipfel, C. (2016). Regulation of pattern recognition receptor signalling in plants. Nature Review Immunology 16: 537–52.

2. Ronald, P.C., and Beutler, B. (2010). Plant and animal sensors of conserved microbial signatures. Science 330: 1061–1064.

3. Spoel, S.H., and Dong, X. (2012). How do plants achieve immunity? Defence without specialized immune cells. Nature Review Immunology 12: 89–100.

4. Boller, T., and Felix, G. (2009). A renaissance of elicitors: perception of microbe-associated molecular patterns and danger signals by pattern-recognition receptors. Annual Review of Plant Biology 60: 379–406.

5. Zhang, J., Li, W., Xiang, T., Liu, Z., Laluk, K., Ding, X., Zou, Y., Gao, M., Zhang, X., Chen, S., Mengiste, T., Zhang, Y., and Zhou, J.M. (2010). Receptor-like cytoplasmic kinases integrate signaling from multiple plant immune receptors and are targeted by a Pseudomonas syringae effector. Cell Host Microbe. 7: 290–301.

6. Thatcher, L., Anderson, J., and Singh, K. (2005). Plant defence responses: what have we learnt from arabidopsis. Functional plant biology 32: 1–19.

7. Ulker, B., and Somssich, I.E. (2005). WRKY transcription factors: from DNA binding towards biological function. Current Opinion in Plant Biology 7: 491–8.

8. Zipfel, C., Kunze, G., Chinchilla, D., Caniard, A., Jones, J.D., Boller, T., and Felix, G. (2006). Perception of the bacterial PAMP EF-Tu by the receptor EFR restricts Agrobacterium-mediated transformation. Cell 125: 749–760.

9. Dangl, J.L., Horvath, D.M., and Staskawicz, B.J., (2013). Pivoting the plant immune system from dissection to deployment. Science 341: 746–51.

10. Sarris, P.F., Cevik, V., Dagdas, G., Jones, J.D., and Krasileva, K.V. (2016). Comparative analysis of plant immune receptor architectures uncovers host proteins likely targeted by pathogens. BMC Biology 19: 8.

11. Zipfel, C., and Felix, G. (2005). Plants and animals: a different taste for microbes? Current Opinion in Plant Biology 8: 353–360.

12. Macho, A.P., and Zipfel, C. (2014). Plant PRRs and the Activation of innate immune Signaling. Molecular Cell 54: 263–272.

13. Newman, M.A., Sundelin, T., Nielsen, J.T., and Erbs, G. (2013). MAMP (microbe-associated molecular pattern) triggered immunity in plants. Front Plant Science. 16: 139.

14. Bahar, O., Mordukhovich, G., Luu, D.D., Schwessinger, B., Daudi, A., Jehle, A.K., Felix, G., and Ronald, P.C. (2016). Bacterial outer membrane vesicles induce plant immune responses. Molecular Plant Microbe Interactions 29: 374–84.

15. Niehl, A., Wyrsch, I., Boller, T., and Heinlein, M. (2016). Double-stranded RNAs induce a pattern-triggered immune signaling pathway in plants. New Phytologist 211: 1008–19.

16. Park, Y.S., Lee, B., and Ryu, C.M. (2016). Getting to PTI of bacterial RNAs: Triggering plant innate immunity by extracellular RNAs from bacteria. Plant Signal Behavior. 11: e1198866.

17. Trdá, L., Boutrot, F., Claverie, J., Brulé, D., Dorey, S. and Poinssot, B. (2015). Perception of pathogenic or beneficial bacteria and their evasion of host immunity: pattern recognition receptors in the front line. Frontiers in Plant Science. doi: 10.3389/fpls.2015.00219

18. Ausubel, F.M. (2005). Are innate immune signaling pathways in plants and animals conserved? Nature Immunology 6: 973–9.

19. Greeff, C., Roux, M., Mundy, J., and Petersen, M. (2012). Receptor-like kinase complexes in plant innate immunity. Frontiers in plant science 3: 209–2012.

a. Shiu SH, Bleecker AB. Plant receptor-like kinase gene family: diversity, function, and signaling. Sci STKE. 2001 Dec 18;2001(113):re22. doi: 10.1126/stke.2001.113.re22. PMID: 11752632.

b. Shiu, S.H:, and Bleecker, A.B. (2001b). Receptor-like kinases from Arabidopsis form a monophyletic gene family related to animal receptor kinases. Proc. Natl. Acad. Sci. USA 98:10763–10768

20. Gómez-Gómez L, Boller T. FLS2: an LRR receptor-like kinase involved in the perception of the bacterial elicitor flagellin in Arabidopsis. Mol Cell. 2000 Jun;5(6):1003–11. doi: 10.1016/s1097-2765(00)80265-8. PMID: 10911994.

21. Li J, Chory J. A putative leucine-rich repeat receptor kinase involved in brassinosteroid signal transduction. Cell. 1997 Sep 5;90(5):929–38. doi: 10.1016/s0092-8674(00)80357-8. PMID: 9298904.

22. Clark SE: Organ formation at the vegetative shoot meristem. Plant Cell 1997, 9:1067–1076.

23. Hecht V, Vielle-Calzada JP, Hartog MV, Schmidt ED, Boutilier K, Grossniklaus U, de Vries SC. The Arabidopsis SOMATIC EMBRYOGENESIS RECEPTOR KINASE 1 gene is expressed in developing ovules and embryos and enhances embryogenic competence in culture. Plant Physiol. 2001 Nov;127(3):803-16. Erratum in: Plant Physiol 2002 Jan;128(1):314. PMID: 11706164; PMCID: PMC129253.

24. Aan den Toorn M, Albrecht C, de Vries S. On the Origin of SERKs: Bioinformatics Analysis of the Somatic Embryogenesis Receptor Kinases. Mol Plant. 2015 May;8(5):762–82. doi: 10.1016/j.molp.2015.03.015. Epub 2015 Apr 9. PMID: 25864910.

25. Sakamoto T, Deguchi M, Brustolini OJ, Santos AA, Silva FF, Fontes EP. The tomato RLK superfamily: phylogeny and functional predictions about the role of the LRRII-RLK subfamily in antiviral defense. BMC Plant Biol. 2012 Dec 2;12:229. doi: 10.1186/1471-2229-12-229. PMID: 23198823; PMCID: PMC3552996.

26. Li B, Ferreira MA, Huang M, Camargos LF, Yu X, Teixeira RM, Carpinetti PA, Mendes GC, Gouveia-Mageste BC, Liu C, Pontes CSL, Brustolini OJB, Martins LGC, Melo BP, Duarte CEM, Shan L, He P, Fontes EPB. The receptor-like kinase NIK1 targets FLS2/BAK1 immune complex and inversely modulates antiviral and antibacterial immunity. Nat Commun. 2019 Nov 1;10(1):4996. doi: 10.1038/s41467-019-12847-6. PMID: 31676803; PMCID: PMC6825196.

27. Hančinský R, Mihálik D, Mrkvová M, Candresse T, Glasa M. Plant Viruses Infecting Solanaceae Family Members in the Cultivated and Wild Environments: A Review. Plants (Basel). 2020 May 25;9(5):667. doi: 10.3390/plants9050667. PMID: 32466094; PMCID: PMC7284659.

28. Safaeizadeh M, Saidi A, Palukaitis P. Molecular characterization of cucumber mosaic virus isolates infecting tomato in Hamedan and Tehran provinces of Iran. Acta Virol. 2015 Jun;59(2):174–8. doi: 10.4149/av_2015_02_174. PMID: 26104334.

29. Chinchilla D, Shan L, He P, de Vries S, Kemmerling B. One for all: the receptor-associated kinase BAK1. Trends Plant Sci. 2009 Oct;14(10):535–41. doi: 10.1016/j.tplants.2009.08.002. Epub 2009 Sep 10. PMID: 19748302; PMCID: PMC4391746.

30. Hančinský R, Mihálik D, Mrkvová M, Candresse T, Glasa M. Plant Viruses Infecting Solanaceae Family Members in the Cultivated and Wild Environments: A Review. Plants (Basel). 2020 May 25;9(5):667. doi: 10.3390/plants9050667. PMID: 32466094; PMCID: PMC7284659.

31. Safaeizadeh M, Saidi A, Palukaitis P. Molecular characterization of cucumber mosaic virus isolates infecting tomato in Hamedan and Tehran provinces of Iran. Acta Virol. 2015 Jun;59(2):174–8. doi: 10.4149/av_2015_02_174. PMID: 26104334.

33. Voloudakis A.E., Mukherjee S.K., Roy A. (2021) Novel Technologies for Transgenic Management for Plant Virus Resistance. In: Sarmah B.K., Borah B.K. (eds) Genome Engineering for Crop Improvement. Concepts and Strategies in Plant Sciences. Springer, Cham. https://doi.org/10.1007/978-3-030-63372-1_7

34. Brustolini, O.J.B., Machado, J.P.B., Condori-Apfata, J.A., Coco, D., Deguchi, M., Loriato, V.A.P., Pereira, W.A., Alfenas-Zerbini, P., Zerbini, F.M., Inoue-Nagata, A.K., Santos, A.A., Chory, J., Silva, F.F., and Fontes, E.P.B. (2015). Sustained NIK-mediated antiviral signalling confers broad spectrum tolerance to begomoviruses in cultivated plants. Plant Biotechnol J. 13:1300–1311

35. Hosseini, S., Schmidt, E.D.L., and Bakker, F.T. (2020). Leucine-rich receptor-like kinase II phylogenetics reveals five main clades throughout the plant kingdom. The Plant Journal 103:547–560

36. Santos, A.A., Carvalhi, C.M., Florentino, L.H., Ramos, H.J.O., Fontes, E.P.B. (2009). Conserved threonine residues within the A loop of the receptor NIK1 differentially regulate the kinase function required for antiviral signaling. PLoS ONE 2009, 4, e5781.

37. Zorzatto C, Machado JPB, Lopes KGV, Nascimento KJT, Pereira WA, Brustolini OJ, Reis PA, Calil IP, Deguchi M, Sachetto-Martins G et al. (2015). NIK1-mediated translation suppression functions as a plant antiviral immunity mechanism. Nature 520:679–682

38. Pyott D.E., Sheehan E., and Molnar A. (2016). Engineering of CRISPR/Cas9-mediated potyvirus resistance in transgene-free Arabidopsis plants. Mol Plant Pathol. 17:1276–88

39. Chandrasekaran J, Brumin M, Wolf D, Leibman D, Klap C, Pearlsman M, Sherman A, Arazi T and Gal-On A. (2016). Development of broad virus resistance in non-transgenic cucumber using CRISPR/Cas9 technology. Mol Plant Pathol. 17: 1140–53. doi: 10.1111/mpp.12375. Epub 2016 Apr 21.

40. J. Li, J. Wen, K.A. Lease, J.T. Doke, F.E. Tax, J.C. Walker (2002). BAK1, an Arabidopsis LRR receptor-like protein kinase, interacts with BRI1 and modulates brassinosteroid signaling. Cell, 110 pp. 213–222

41. K.H. Nam, J. Li (2002). BRI1/BAK1, a receptor kinase pair mediating brassinosteroid signaling. Cell, 110 pp. 203–212

42. D. Chinchilla, C. Zipfel, S. Robatzek, B. Kemmerling, T. Nurnberger, J.D. Jones, G. Felix, T. Boller (2007). A flagellin-induced complex of the receptor FLS2 and BAK1 initiates plant defence Nature, 448 pp. 497–500

37b. Carvalho CM, Santos AA, Pires SR, Rocha CS, Saraiva DI, Machado JP, Mattos EC, Fietto LG, Fontes EPB. Regulated nuclear trafficking of rpL10A mediated by NIK1 represents a defense strategy of plant cells against virus. PLoS Pathog. 2008c; 4:e1000247. [PubMed: 19112492]

38. Santos AA, Lopes KVG, Apfta JAC, Fontes EPB (2010). NSP-interacting kinase, NIK: a transducer of plant defence signalling. J. Exp. Bot. 61:3839–3845. [PubMed: 20624762]

